# Characterization of the signalling modalities of prostaglandin E2 receptors EP2 and EP4 reveals crosstalk and a role for microtubules

**DOI:** 10.1101/2020.10.06.328351

**Authors:** Ward Vleeshouwers, Koen van den Dries, Sandra de Keijzer, Ben Joosten, Diane S. Lidke, Alessandra Cambi

## Abstract

Prostaglandin E2 (PGE2) is a lipid mediator that modulates the function of myeloid immune cells such as macrophages and dendritic cells (DCs) through the activation of the G protein-coupled receptors EP2 and EP4. While both EP2 and EP4 signalling leads to an elevation of intracellular cyclic adenosine monophosphate (cAMP) levels through the stimulating Gα_s_ protein, EP4 also couples to the inhibitory Gα_i_ protein to decrease the production of cAMP. The receptor-specific contributions to downstream immune modulatory functions are still poorly defined. Here, we employed quantitative imaging methods to characterize the early EP2 and EP4 signalling events in myeloid cells and their contribution to the dissolution of adhesion structures called podosomes, which is a first and essential step in DC maturation. We first show that podosome loss in DCs is primarily mediated by EP4. Next, we demonstrate that EP2 and EP4 signalling leads to distinct cAMP production profiles, with EP4 inducing a transient cAMP response and EP2 inducing a sustained cAMP response only at high PGE2 levels. We further find that simultaneous EP2 and EP4 stimulation attenuates cAMP production, suggesting a reciprocal control of EP2 and EP4 signaling. Finally, we demonstrate that efficient signaling of both EP2 and EP4 relies on an intact microtubule network. Together, these results enhance our understanding of early EP2 and EP4 signalling in myeloid cells. Considering that modulation of PGE2 signalling is regarded as an important therapeutic possibility in anti-tumour immunotherapy, our findings may facilitate the development of efficient and specific immune modulators of PGE2 receptors.

## Introduction

The ability of cells to respond to their environment is critical for their function. Important players for transmitting extracellular information into intracellular signalling events are the G protein-coupled receptors (GPCRs) [1]. The spatiotemporal organization of GPCRs within the cell membrane allows these receptors to elicit fine-tuned cellular responses to different ligands.

Prostaglandins are lipid mediators that represent an abundant type of GPCR ligand. Prostaglandins are derived from cyclooxygenase (COX)-catalyzed metabolism of arachidonic acid and exhibit versatile actions in a wide variety of tissues [2; 3]. Prostaglandin E2 (PGE2) signals via the four GPCRs EP1-4, expressed in various combinations at the plasma membrane of cells [4](REF). PGE2 modulates several key immunological processes including the activation, migration and cytokine production of different immune cells such as dendritic cells (DCs), macrophages and T lymphocytes [3; 5; 6; 7; 8]. Despite being a known mediator of inflammation, increased PGE2 concentrations have been associated with a highly immunosuppressive tumor microenvironment (TME) of several cancer types [9; 10; 11; 12; 13].

DCs are commonly observed in the TME of solid tumors [14]. Yet, despite their potential to generate anti-tumor immunity, TME-resident DCs often exhibit impaired or defective function [15]. The high PGE2 levels in the TME might play a role since PGE2 promotes IL-10 production by DCs [16]. On the other hand, PGE2 is also important for inducing the highly migratory phenotype typical of mature DCs and which is crucial in immunity [6]. Understanding how PGE2 exerts its dual function in DCs can offer novel leads to reverse unwanted DC immunosuppression in the context of anti-tumor immunity.

PGE2 modulates DC function exclusively via EP2 and EP4 [6; 17; 18]. For example, PGE2 has previously been shown to induce the dissolution of podosomes, which are actin-rich adhesion structures involved in tissue-resident immature DC migration, through the cAMP-PKA-RhoA signaling axis downstream of EP2 and EP4 [8]. PGE2-induced podosome dissolution is an important step towards DC maturation, but the receptor-specific contributions to these processes are still poorly defined.

Signaling via EP2 and EP4 is predominantly transduced by the stimulating Gα protein (Gα_s_), leading to increased activity of adenylate cyclase (AC) and subsequent elevation of intracellular cyclic adenosine monophosphate (cAMP) levels [19; 20]. An important difference between EP2 and EP4 is the reported capacity of EP4 to also couple to inhibitory Gα protein (Gα_i_), thereby inhibiting cAMP formation and activating a phosphatidylinositol 3-kinase (PI3K) pathway [21; 22]. Furthermore, in contrast to EP2, EP4 is rapidly internalized upon ligand binding [23; 24; 25]. Altogether, these observations suggest that signal modalities (intensity, duration, downstream effectors) likely differ between EP2 and EP4 and a better understanding of EP2 and EP4 signalling modalities is key to understand PGE2 effects in DC biology.

Here, we aimed to characterize EP2 and EP4 early signaling events in response to PGE2 in myeloid cells. We first demonstrate that in DCs, PGE2 leads to podosome dissolution primarily through EP4 signalling. Next, we show that selective EP2 and EP4 stimulation leads to distinct cAMP production profiles and suggest reciprocal control of receptor signalling efficiency. Finally, we demonstrate that the integrity of the cortical microtubule network is important for efficient EP2 and EP4 signalling. Modulation of PGE2 signalling is considered an important therapeutic possibility in anti-tumour immunotherapy. Our findings enhance our understanding of early EP2 and EP4 signaling and may thereby facilitate the development of efficient and specific modulators of PGE2 signalling receptors that can contribute to reverse tumor immunosuppression [26].

## Materials and methods

### Chemicals and reagents

Cells were treated with several compounds that activated or inhibited EP2 and EP4: EP2 agonist (R)-Butaprost (Sigma), EP4 agonist L-902688 (Cayman Chemicals), EP2 antagonist AH6809 (Cayman Chemicals), EP4 antagonist GW627368X (Cayman Chemicals) or AH23848 (Cayman Chemicals), pertussis toxin (TOCRIS biosciences), PGE2 (Cayman Chemicals), Pertussis Toxin (PTx, Calbiochem, San Diego, CA) and nocodazole (Sigma). Compounds used for immunofluorescence staining were mouse anti-vinculin antibody (Sigma, V9131), Goat anti-Mouse-(H&L)-Alexa488 and Goat anti-Mouse-(H&L)-Alexa647 secondary antibodies (Invitrogen), Alexa488-conjugated phalloidin (Invitrogen, A12379) and Texas Red-conjugated phalloidin (Invitrogen, T7471), Mowiol (Sigma).

### Cell culture

RAW 246.7 cells were cultured in RPMI-1640 medium (Gibco) supplemented with 10% Fetal Bovine Serum (FBS, Greiner Bio-one), 1mM Ultra-glutamine (BioWitthaker) and 0.5% Antibiotic-Antimytotic (AA, Gibco). iDCs were derived from PBMCs as described previously [27; 28] and cultured in RPMI 1640 medium (Gibco) supplied with 10% Fetal Bovine Serum (FBS, Greiner Bio-one). Transfections with t-Epac-vv [29] (gift from K. Jalink), Gα_s_-GFP (gift from M. Rasenick), Gα_i_-GFP and Gα_i1_-Citrine [30] (gift from A. Gilman), Gγ_2_-CFP and Gβ_1_ wildtype (both gifts from M. Adjobo-Hermans) were performed with Fugene HD (Roche) according to the manufacturer protocol and imaged after 24 h. Stable cell lines expressing Gα_s_-GFP and Gα_i_-GFP was maintained using the appropriate antibiotics. Cells were plated one day prior to measurements or transfection in Willco dishes (Willco Wells BV) at 400.000 cells/dish or in 96 well-plate (microplate BD Falcon) at 40.000 cells/well or in 4-well Lab-Tek II chambered coverglass (Nunc) at 100.000 cells/chamber. Prior to imaging, the medium was replaced with 1 ml RPMI medium without phenol red to avoid background fluorescence.

### Podosome dissolution assay and widefield immunofluorescence

For agonist experiments, iDCs were treated with (R)-Butaprost, L-902688 or 10 μM PGE2 for 10 min. For antagonist and pertussis toxin experiments, iDCs were pretreated with 3 μM AH6809 for 1 h, 10 μM GW627368X for 1 h, 100 ng/ml pertussis toxin for 16 hrs as previously described [22] or left untreated prior to the addition of PGE2. After stimulation, iDCs were fixed in 3.7% (w/v) formaldehyde in PBS for 10 min. Cells were permeabilized in 0.1% (v/v) Triton X-100 in PBS for 5 min and blocked with 2% (w/v) BSA in PBS. The cells were incubated with mouse anti-vinculin antibody for 1 h. Subsequently, the cells were washed with PBS and incubated with GaM-(H&L) secondary antibody and phalloidin for 45 min. Lastly, samples were washed with PB prior to embedding in Mowiol. Cells were imaged on a Leica DM fluorescence microscope with a 63× PL APO 1.3 NA oil immersion lens and a COHU high-performance integrating CCD camera (COHU, San Diego, CA) or a Zeiss LSM 510 microscope equipped with a PlanApochromatic 63x/1.4 NA oil immersion objective. Images were analyzed using Fiji-based software [31].

### FRET experiments

RAW macrophages expressing t-Epac-vv were imaged using a BD Pathway high-content imaging inverted widefield microscope (BD biosciences) equipped with a 20X 0.75 N.A. objective (Olympus LUCPLFLN). A mercury metal halide lamp combined with an excitation filter (440/10) was used to excite mTurqoise. The fluorescence emission was filtered using a dichroic mirror (458-DiO1) and filters (479/40 and 542/27 for mTurquoise and Venus emission, respectively). Emission was collected by a high-resolution cooled CCD camera (1344×1024 pix, 0.32 μm/pix). Samples were prepared in a 96 well-plate (microplate BD Falcon) from which the inner 60 wells were used. Cells were pretreated with with 100 ng/ml pertussis toxin for 16 hrs or left untreated before adding 3 μM AH6809 for 1 h, or 10 μM GW627368X for 1 h, with and without 5 μM nocodazole for 20 min,. Six mTurquoise and Venus emission images were acquired followed by automated addition of PGE2 and subsequent acquisition of another 20 mTurquoise and Venus emission images (t_lag_=10 s). The mean fluorescence intensity of the Venus and mTurqoise signal in a cell was corrected by subtraction of the background signal in each image and channel before dividing the Venus over mTurqoise mean fluorescence intensity to obtain the FRET ratio. Values were normalized to the average ratio value of the first six prestimulus data points.

### FLIM experiments

Frequency-domain FLIM experiments on transfected RAW macrophages were performed using a Nikon TE2000-U inverted widefield microscope and a Lambert Instruments Fluorescence Attachment (LIFA; Lambert Instruments) for lifetime imaging. A light-emitting diode (Lumiled LUXEON III, λ_max_ = 443 nm) modulated at 40 MHz was used to excite CFP. Fluorescence detection was performed by a combination of a modulated (40 MHz) image intensifier (II18MD; Lambert Instruments) and a 640×512 pixel CCD camera (CCD-1300QD; VDS Vosskühler). The emission of CFP was detected through a narrow emission filter (475/20 nm; Semrock) to suppress any fluorescence emission from the Citrine fluorophore. FLIM measurements were calibrated with a 1 μM solution of pyranine (HPTS), the lifetime of which was set to 5.7 ns. All FLIM images were calculated from phase stacks of 12 recorded images, with exposure times of individual images ranging from 200 to 400 ms. A USH-102DH 100 W mercury lamp (Nikon) was used for acceptor photobleaching. Cells were pretreated with 25 μM AH23848 for 1 h or left untreated and cells were stimulated with 10 μM PGE2 or 10 μM Butaprost.

## Results

### EP4 primarily contributes to PGE2-induced podosome dissolution in DCs

To assess different contributions of EP2 and EP4 in mediating PGE2 signalling in DCs, we determined the effect of receptor-specific inhibition or stimulation in podosome dissolution. We first treated immature DCs (iDCs) with PGE2 or selective EP2 and EP4 agonists and quantified the number of podosomes per cell (**Figure 1A,B**). In line with our previous observations, addition of PGE2 resulted in an almost complete loss of podosomes in iDCs. Interestingly, both EP2- and EP4-specific stimulation reduced the number of podosomes, with EP4 agonist stimulation being slightly more efficient (**Figure 1B**). These results indicate that individual EP2 and EP4 downstream signalling can lead to podosome dissolution.

**Figure 1.**
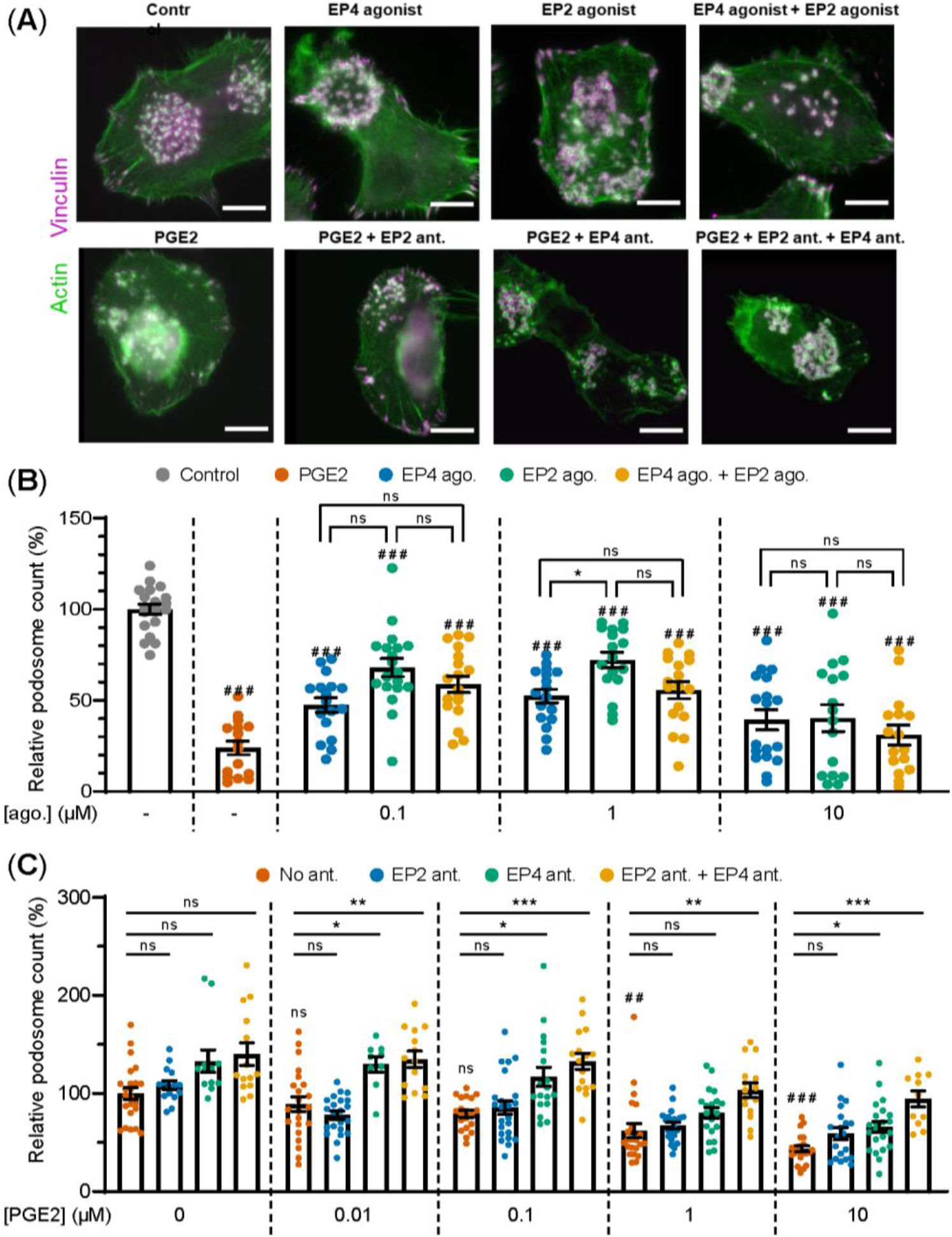
PGE2-induced podosome dissolution in human iDCs is mostly mediated by EP4. (**A**) Representative images of PBMC-derived iDCs that were left untreated or were treated with 1 μM EP4 agonist L-902688, 1 μM EP2 agonist (R)-Butaprost, both 1 μM L-902688 and 1 μM (R)-Butaprost, 1 μM PGE2 alone or 1 μM PGE2 after pretreatment with EP2 antagonist (ant.) AH6809, EP4 antagonist GW627368X or both AH6809 and GW627368X. Cells were stained for actin (green) and vinculin (magenta). Scale bar = 10 μm. Scale bar = 10 μm. (**B**) iDCs were treated with different concentrations of EP2 agonist (ago.) (R)-Butaprost, EP4 agonist L-902688 or both (R)-Butaprost and L-902688. Cells were stained for actin and vinculin and the number of podosomes per image was quantified and normalized to untreated control. Cells treated with 10 μM PGE2 were included as positive control. The error bars represent mean ± SEM. Data presented are from 2 different donors. ns = not significant, *P<0.05; ^###^P<0.001 versus untreated control, Welch ANOVA with Dunnett’s T3 multiple comparison test. (**C**) iDCs were treated with different concentrations of PGE2 with or without pretreatment with EP2 antagonist (ant.) AH6809, EP4 antagonist GW627368X or both AH6809 and GW627368X. Cells were stained for actin and vinculin and the number of podosomes per image was quantified and normalized to untreated control. The error bars represents mean ± SEM. Data presented are from three different donors. ns = not significant, *P<0.05, **P<0.01, ***P<0.001; ^##^P<0.01, ^###^P<0.001 versus untreated control, Welch ANOVA with Dunnett’s T3 multiple comparison test.

To better investigate the respective contribution of EP2 and EP4 signalling after the addition of their natural ligand PGE2, we pretreated the cells with selective EP2 and EP4 antagonists before PGE2 addition and subsequently quantified podosome dissolution. **Figure 1C** shows that inhibition of EP4 attenuates podosome dissolution upon stimulation with 0.01-0.1 μM PGE2, while blocking of EP2 has no effect. This indicates that at lower PGE2 concentrations, EP4 is responsible for the induction of podosome loss. Interestingly, at PGE2 concentrations ≥ 1 μM, EP4 blocking attenuates podosome dissolution only when EP2 antagonist is co-administered, suggesting that EP2 triggering by PGE2 could somehow influence EP4 activity.

These results show that the use of EP agonists does not allow for the detection of the differential contribution of the receptors in mediating PGE2 signalling. Therefore, the use of selective receptor antagonists in combination with the natural ligand PGE2 was chosen to define the individual contributions of EP2 and EP4 in subsequent experiments.

### EP2 and EP4 differentially stimulate cAMP production

PGE2-induced podosome loss in DCs is mediated by the cAMP-PKA-RhoA signaling axis downstream of EP2 and EP4 [8].Since our results strongly suggest that EP4 is primarily responsible for podosome loss, we sought to determine whether EP4 induces stronger cAMP responses to PGE2 than EP2. To determine the individual contribution of EP2 and EP4 to the PGE2-induced increase of intracellular cAMP levels, we measured the onset of cAMP production in living RAW macrophages, which endogenously express both EP2 and EP4 [32], using ratio measurements of the Förster Resonance Energy Transfer (FRET)-based cAMP sensor t-Epac-vv [29]. Since the binding of cAMP to t-Epac-vv reduces FRET between the mTurquoise donor and Venus acceptor fluorophores, a decreased FRET ratio in the macrophages is a direct measure of cAMP production (**Figure 2A,B**). After the addition of PGE2, cAMP was produced immediately and reached a maximum concentration after about 40 seconds, subsiding to lower levels after 200 seconds (**Figure 2C**). To compare the cAMP kinetics across difference treatment conditions, we quantified the peak of cAMP production and the production rate, as shown in **Figure 2D**. Both parameters scaled with increasing PGE2 concentrations, indicating that the rate and the magnitude of the induced cAMP response is dose-dependent (**Figure 2E**).

**Figure 2.**
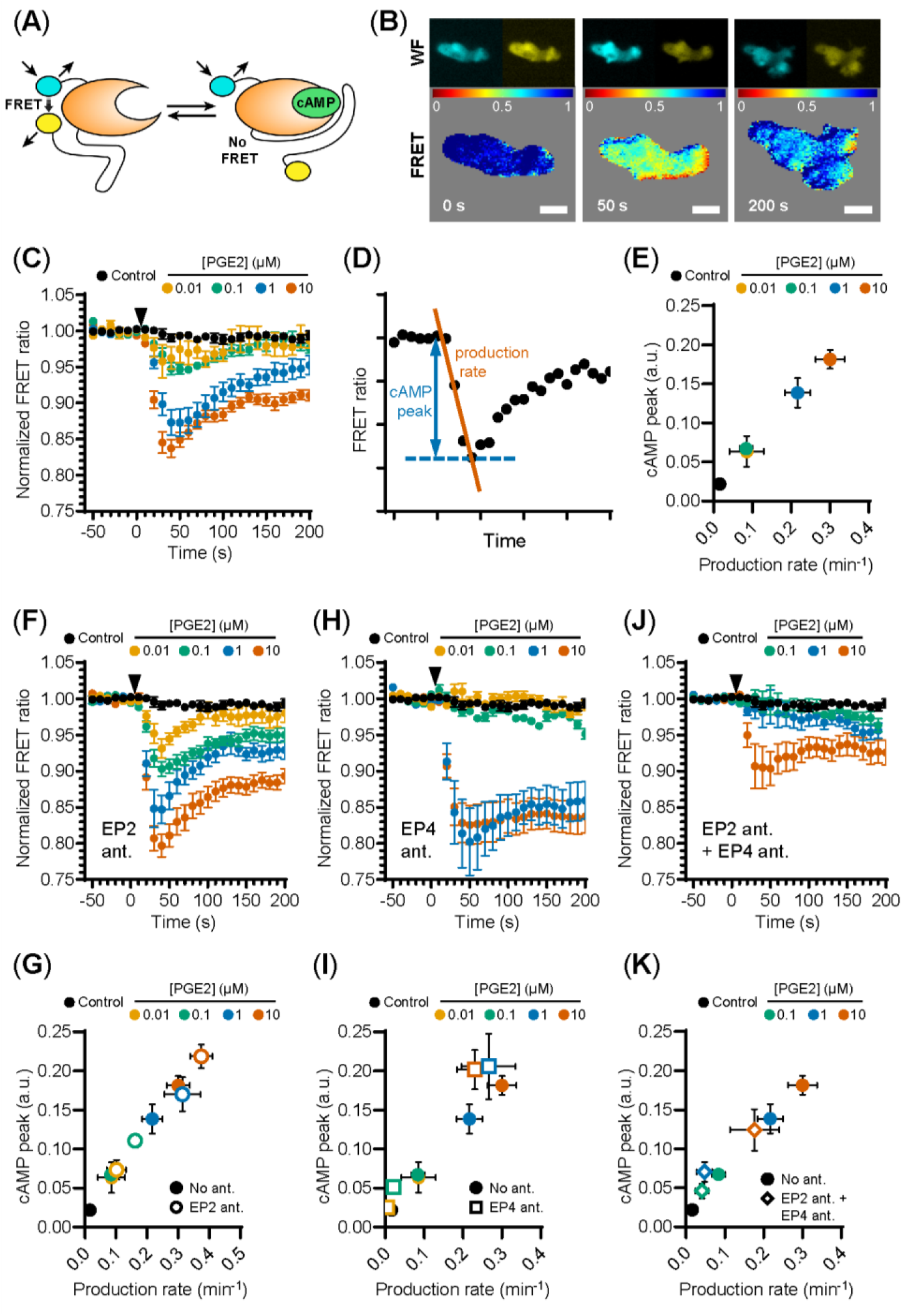
EP2 and EP4 induce distinct cAMP responses. (**A**) Schematic illustration of intramolecular cAMP FRET sensor t-Epac-vv. Binding of cAMP to t-Epac-vv reduces FRET between the mTurquoise donor and Venus acceptor fluorophores of t-Epac-vv, making a decreased ratio of the fluorescent intensities a direct measure of cAMP accumulation (adapted from (REF)). (**B**) The mTurquoise (cyan) and Venus (yellow) signal were acquired with widefield microscopy (WF panels). After background subtraction in each image and channel, the FRET ratio was calculated as the Venus intensity over the mTurqoise intensity for each timepoint and was normalized to the average of prestimulus values (FRET panels). Normalized FRET values range from 0 (red) to 1 (blue). Scale bar = 5 μm. (**C**) FRET ratios of t-Epac-vv before and after the addition of different PGE2 concentrations were measured in transiently transfected RAW macrophages. A control was performed with the addition of buffer only. The data presented are mean ± SEM from ≥5 cells per condition. (**D**) Example FRET curve that illustrates the definition of the relative cAMP peak and cAMP production rate. The amplitude of the cAMP peak was defined as the maximal decrease in FRET ratio. The cAMP production rate was quantified by determining the slope between the final prestimulus timepoint and the timepoint at which minimal FRET ratios were observed using a linear fit over all included timepoints. (**E**) The cAMP production peak and the cAMP production rate were measured from the FRET curve of individuals cells from (**C**) and the average peak was plotted as a function of the average production rate per condition. The error bars represent SEM for both parameters. (**F, H, J**) FRET ratios were measured after the addition of PGE2 in cells pretreated with EP4 antagonist (ant.) GW627368X (**F**), pretreated with EP2 antagonist AH6809 (**H**) or pretreated with both GW627368X and AH6809 (**J**). The data presented are mean ± SEM from ≥4 cells per condition. (**G, I, K**) The relative cAMP production peak and the cAMP production rate were measured from (**F**), (**H**) and (**J**), respectively. The error bars represent SEM for both parameters.

Compared to PGE2 only, EP2 inhibition led to higher cAMP levels at all tested PGE2 concentrations, while cAMP concentrations subsided to a similar extent (**Figure 2F**). The PGE2-induced cAMP production rate and cAMP peak remained dose-dependent upon EP2 inhibition as both parameters scaled with PGE2 concentration (**Figure 2G**). These results indicate that EP2 blockade increases the signaling efficiency of EP4 in response to PGE2. Inhibition of EP4 led to dramatically different cAMP production. In contrast to EP2 inhibition, robust cAMP production was not observed until 1 μM PGE2 when EP4 signaling was blocked (**Figure 2H,I**). Furthermore, this strong cAMP response did not attenuate as observed in the absence of EP4 inhibition. Compared to PGE2 only, the magnitude of the strong cAMP response observed upon EP4 inhibition suggests that EP4 response did not attenuate as observed in the absence of EP4 inhibition. Compared to PGE2 only, the magnitude of the strong cAMP response observed upon EP4 inhibition suggests that EP4 activity may somehow impair the signaling efficiency of EP2. To ascertain that EP2 and EP4 are completely blocked by the antagonist concentrations used in our experiments, we measured cAMP production upon simultaneous inhibition of EP2 and EP4 (**Figure 2J**). Pretreatment with both antagonists effectively inhibited total cAMP production at 0.1 and 1 μM PGE2, showing that both receptors are completely blocked at physiological concentrations of PGE2 (**Figure 2J,K**).

Our results demonstrate that the selective stimulation of EP2 and EP4 by PGE2 induces kinetically distinct cAMP production profiles. While PGE2-EP4 signalling results in a fast and transient cAMP production that linearly increases with increasing ligand concentrations, PGE2-EP2 signalling is induced only by PGE2 concentrations above 1 μM and cAMP production and is more prolonged. We also show that co-stimulation of EP2 and EP4 mutually dampens their signaling efficiency, as both receptors induce higher cAMP production when they are individually triggered by PGE2.

### EP4-coupled Gα_**i**_ **finetunes the PGE2-induced cAMP production**

Given that EP2 and EP4 differentially control cAMP dynamics, we sought to identify factors that contribute to these differences. Since the inhibitory G protein Gα_i_ has been shown to couple to EP4 [22], we hypothesized that Gα_i_ dampens the PGE2-induced cAMP response in cells expressing EP4. To demonstrate that EP4 selectively activates Gα_i_ also in macrophages, we performed fluorescence lifetime imaging (FLIM) to measure FRET between cyan fluorescent protein (CFP)-tagged Gγ (Gγ-CFP) and Citrine-tagged Gα_i_ (Gα_i_-Citrine). The fluorescent lifetime of the FRET donor decreased upon co-expression with the acceptor and was restored to control levels upon acceptor photobleaching (**Figure 3A**), indicating that FRET occurred between Gγ-CFP and Gα_i_-Citrine. Since Gα_i_ is known to undergo conformational rearrangements upon activation [33] and FRET between Gγ-CFP and Gα_i_-Citrine is likely affected by such rearrangements, a shift in fluorescence lifetime is expected upon EP4 stimulation. Treatment with PGE2 induced a gradual reduction in the lifetime of the donor fluorophore, whereas no shift in the lifetime phase was observed upon either inhibition of EP4 or selective stimulation of EP2 (**Figure 3B**). These findings confirm that PGE2 induces Gα_i_ activation via EP4 only.

**Figure 3.**
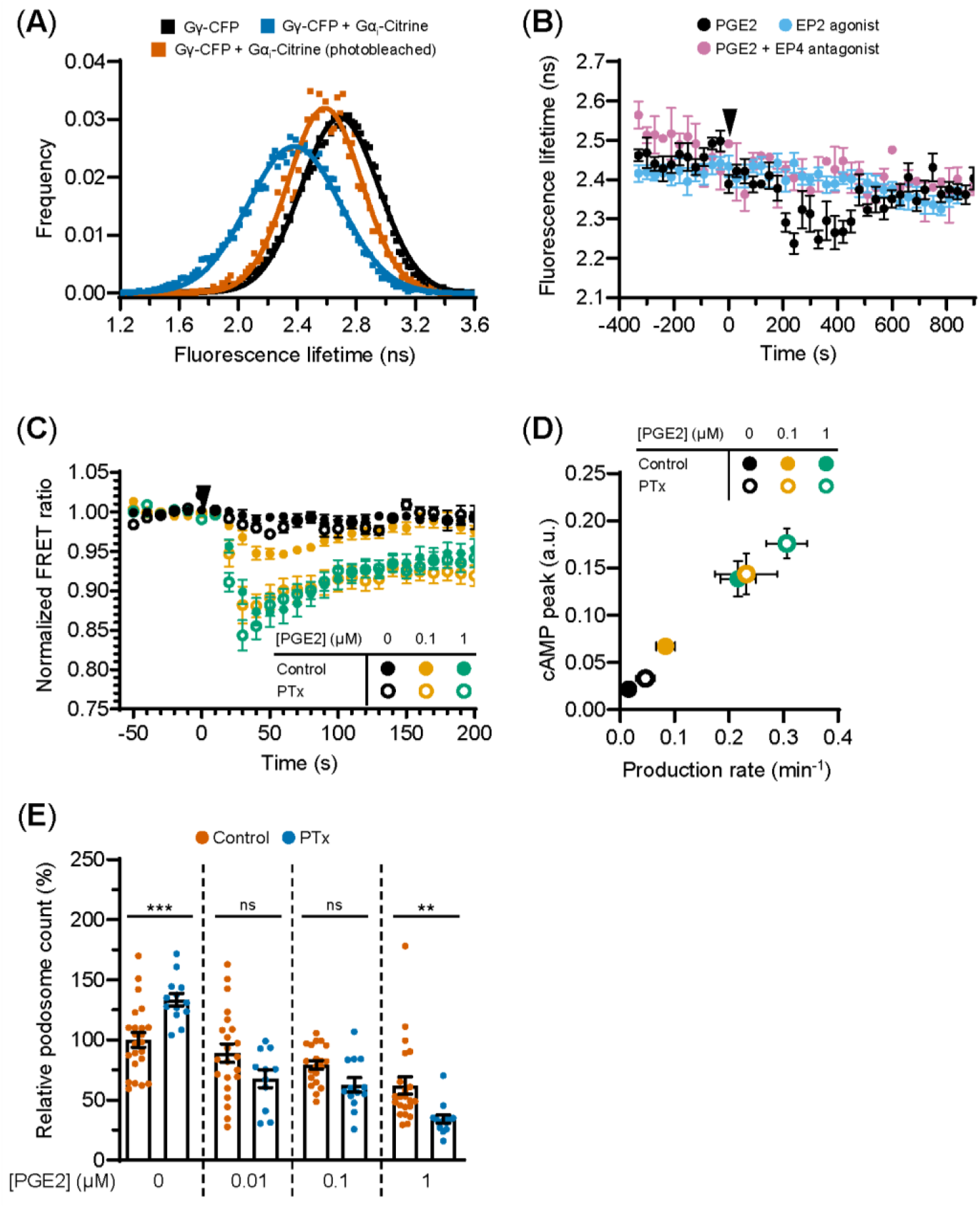
EP4-coupled Gai dampens the PGE2-induced cAMP production. (**A**) RAW macrophages were transfected with Gγ-CFP only or with Gγ-CFP (donor), Gα_i_-Citrine (acceptor) and Gβ wildtype together. The average lifetime of Gγ-CFP for individual cells were calculated from frequency-domain FLIM images and the distributions were fitted with a Gaussian profile (solid lines) to obtain the average lifetimes. Photobleaching of Gα_i_-Citrine was used as a control for the occurrence of FRET. (**B**) The average donor lifetime in cells expressing both donor and acceptor is plotted before and after addition of 10 μM PGE2 in absence or presence of EP4 antagonist AH23848 or after addition of 10 μM EP2 agonist Butaprost. The data presented are mean ± SEM from >5 cells. (**C**) FRET ratios of t-Epac-vv before and after addition of PGE2 were measured in transiently transfected RAW macrophages that were left untreated or were pretreated with Gα_i_ inhibitor pertussis toxin (PTx). Controls were performed with the addition of buffer only. The data presented are mean ± SEM of measurements from ≥4 cells per condition. (**D**) The cAMP peak and the cAMP production rate were quantified as described in Figure 2D from (**C**) and the average peak was plotted as a function of the average production rate per condition. The error bars represent SEM for both parameters. (**E**) iDCs were treated with different concentrations of PGE2 with or without PTx pretreatment. Cells were stained for actin and vinculin and the number of podosomes per image was quantified and normalized to untreated control. The error bars represents mean ± SEM. Data presented are from three different donors. **P<0.01, ***P<0.001, Welch ANOVA with Dunnett’s T3 multiple comparison test.

To determine the consequences of EP4-mediated Gα_i_ activation on PGE2 signaling, we measured cAMP elevation using t-Epac-vv upon inhibition of Gα_i_ with pertussis toxin (PTx). Gα_i_ blockade significantly enhanced the cAMP peak concentrations and production induced by 0.1 μM PGE2 and by 1 μM PGE2, albeit at a lower extent (**Figure 3C**), indicating that Gα_i_ attenuates cAMP production most strongly at lower PGE2 concentrations. The effect of Gα_i_ inhibition on cAMP production is more clearly depicted in Figure 3D, where a higher cAMP peak and an increased production rate are observed after addition of PTx.

Next, to investigate whether EP4-mediated Gα_i_ activation would enhance cAMP-dependent processes such as podosome dissolution, we determined PGE2-mediated podosome loss in iDCs with or without PTx treatment. We found that Gα_i_ inhibition led to slightly increased podosome loss at all PGE2 concentrations tested, with 1 μM PGE2 being statistically significant while 0.01 and 0.1 μM PGE2 show a non-significant but clear trend (**Figure 3E**). It should be considered that such low concentrations of PGE2 are less powerful in inducing podosome dissolution, which means that PTx effect is more difficult to assess. This result indicates that the Gα_i_-mediated dampening of cAMP production also affects cellular decisions downstream of EP2 and EP4.

Together, these findings indicate that Gα_i_ dampens the onset of cAMP production, suggesting that the PGE2-EP4-Gα_i_ axis might act as signalling gatekeeper when low PGE2 levels slightly fluctuate.

### EP2- and EP4-mediated signalling requires cortical microtubule integrity

Since the interplay between G proteins and tubulin is well documented as well as their localization along microtubules [34; 35; 36], we investigated whether microtubule integrity is important for PGE2-induced cAMP production. We found that microtubule disruption deregulates PGE2-induced cAMP elevation (**Figure 4A**). More specifically, when both receptors are activated, attenuation of the cAMP response by nocodazole was only observed at 1 μM PGE2 and not at 0.1 μM PGE2 (**Figure 4A,B**). Upon EP2 inhibition, however, the cAMP production rate and the maximum cAMP levels induced by PGE2-EP4 were reduced at all PGE2 concentrations tested (**Figure 4C,D**). Finally, EP4 inhibition revealed that the PGE2-EP2 strong and sustained cAMP response is completely prevented by microtubule disruption (**Figure 4E,F**). These results demonstrate that the Gα_s_-mediated cAMP response to PGE2 relies on an intact microtubule network and that disruption of this network reduces the signaling efficiency of both EP2 and EP4, with EP2 activity being significantly more sensitive to microtubule integrity than EP4 activity.

**Figure 4.**
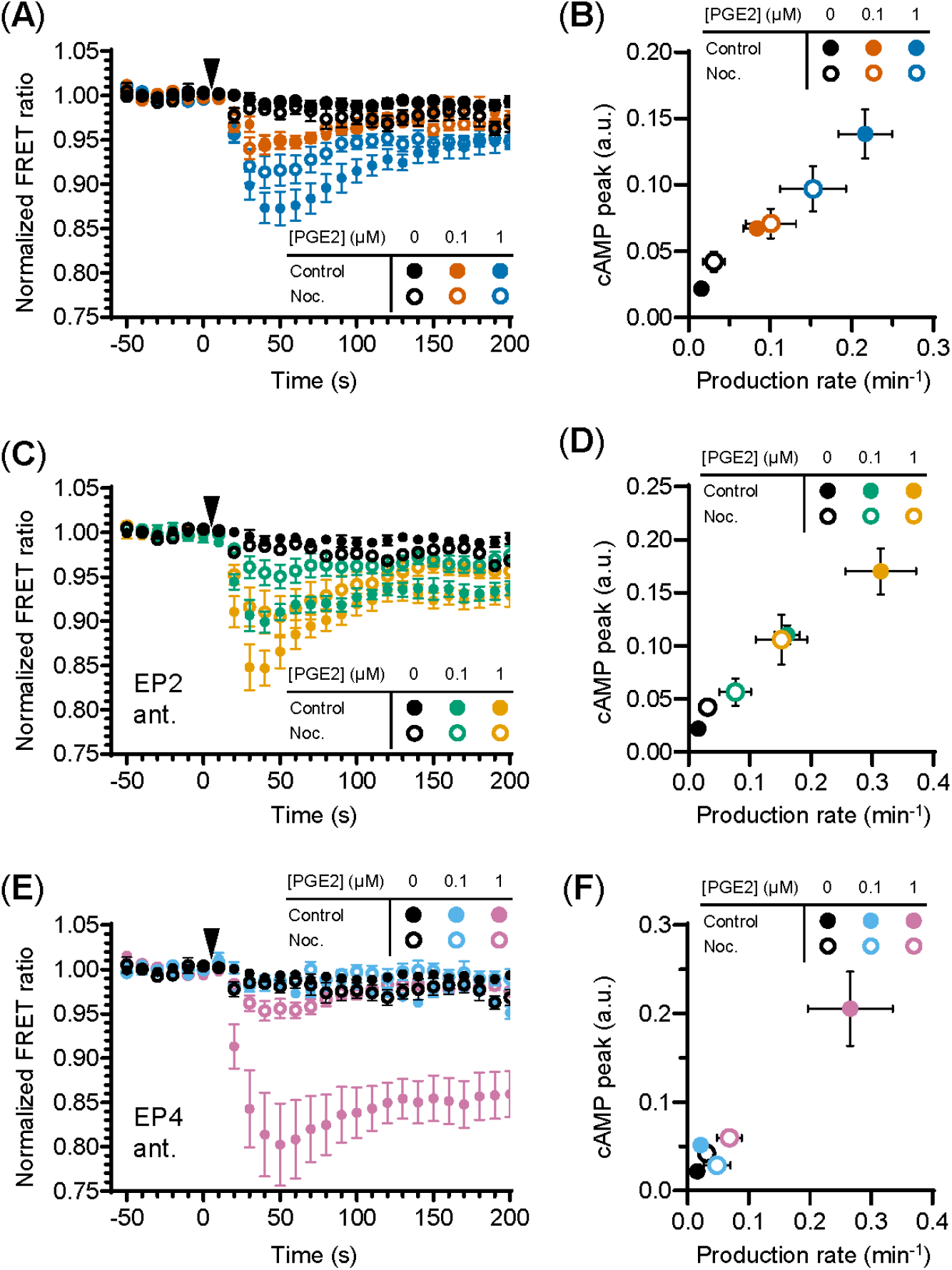
Efficient signaling of EP2 and EP4 relies on microtubule integrity. (**A, C, E**) The FRET ratio of t-Epac-vv was measured in cells that were untreated or pretreated with nocodazole (Noc.) before and after addition of PGE2. Shown are the ratios obtained in cells in the absence of antagonists (**A**), in the presence of EP2 antagonist AH6809 (**C**) or EP4 antagonist GW627368X (**E**). Controls were performed with the addition of buffer only. The data are mean ± SEM from ≥5 cells per condition. (**B, D, F**) The cAMP production peak and the cAMP production rate were measured from the FRET curve of individual cells from (**A**), (**C**) and (**E**), respectively, and the average peak was plotted as a function of the average production rate per condition. The error bars represent SEM for both parameters.

## Discussion

This study characterized the EP2 and EP4 signalling modalities to better understand DC and macrophage responses elicited by PGE2. Our first important observation is that selective activation of EP2 and EP4 by agonists leads to different outcomes compared to activation by PGE2 in the presence of selective receptor antagonists. More specifically, when the receptors are individually activated by a selective agonist, podosome dissolution is almost equally induced by EP2 and EP4, whereas podosome dissolution is mostly mediated by EP4 after the addition of natural ligand PGE2 in the presence of selective antagonists. Throughout this study we consistently applied selective antagonists to determine individual receptor contributions to PGE2 signalling and show that 1) both EP2 and EP4 signal more efficiently when selectively activated by their natural ligand PGE2; 2) EP4 induces dose-dependent and transient cAMP production, whereas EP2 induces a sustained cAMP response only at high PGE2 concentration; 3) EP4-linked Gα_i_ dampens both PGE2-induced cAMP generation and podosome dissolution; 4) microtubule disruption obstructs efficient signaling of both receptors with a very strong effect particularly on EP2 activity.

We here show that also PGE2-induced podosome loss in iDCs [18] is differentially controlled by EP2 and EP4. PGE2-induced podosome dissolution is a first step towards the acquisition of a fast migratory phenotype by DCs [18; 37]. In fact, PGE2 is an important factor to induce DC maturation and by using selective agonists, both EP2 and EP4 have been proposed to play similar roles in this process [6; 16]. Our results rather suggest that this might not be the case and that EP4 is likely the most predominant receptor mediating PGE2 signalling leading to migratory mature DCs. This is in line with previous findings in gene-targeting experiments in mice, where PGE2-EP4 signalling was found to promote migration and maturation of Langerhans cells, thereby initiating skin immune responses [38]. Similarly, other PGE2-mediated immunological processes such cytokine production and T cell activation have been reported to be controlled differently by EP2 and EP4 [39; 40; 41]. Knockdown of EP2 or EP4 in DCs possibly in combination with the use of agonists and antagonists might eventually help to clarify these differences. However, since EP2 and EP4 are always co-expressed in DCs, one will have to rule out that knockdown of one receptor will not affect expression patterns of the other receptor.

Early studies characterizing the EP receptor signaling capacity have mostly used cells that overexpress either EP2 or EP4 [22; 23; 25; 42; 43; 44], which makes it challenging to determine the differential contribution of the receptors when they are co-expressed. Here, we have addressed this question and measured the early onset of cAMP production in cells that endogenously express both EP2 and EP4. Using selective EP2 and EP4 antagonists, we demonstrate that EP2 induces sustained cAMP, whereas EP4-mediated cAMP production is faster but more transient. This difference may be partially explained by the fact that EP4, and not EP2, is internalized shortly after stimulation with PGE2, which halts further signalling [23; 45]. Furthermore, our results showing a sustained EP2-induced cAMP production are in line with the previous observation that EP2 is the main cAMP generator after extended PGE2 stimulation [42]. We also know that EP4 can couple to both Gα_s_ and Gα_i_ [22]. Here, we provide additional evidence that Gα_i_ is only linked to EP4 and not to EP2 and that Gα_i_ attenuates the cAMP response induced by low PGE2 concentrations. Given that several GPCRs do not precouple with Gα_i_ [46], it would be important to determine how and when EP4 and Gα_i_ interact. In a recent study, hidden Markov modeling classified G proteins into four diffusion states, of which the slowest two states represent G proteins that interact in hot spots for GPCR activation [47]. The same study employed single-molecule tracking to show that adrenergic receptors and Gα_i_ proteins interact only transiently within these hot spots [47]. Single-molecule imaging methods are excellent tools to understand the fundamental principles of G protein dynamics and could be exploited to better understand the molecular mechanisms regulating the spatiotemporal interaction between EP4 and Gα_s_ or Gα_i_, which could shape the cAMP production profile.

Our FRET measurements also reveal that the cAMP response of EP4 is dose-dependent, whereas the EP2-induced cAMP production is negligible at low PGE2 concentrations and strong at high PGE2 concentrations. EP4 has a higher affinity for PGE2 than EP2, as indicated by dissociation constants of 0.59 nM and 13 nM, respectively [48]. The high affinity of EP4 explains its responsiveness to low PGE2 concentrations, but the apparent irresponsiveness of EP2 to PGE2 concentrations below 1 μM cannot be explained by its lower affinity for PGE2, based on the magnitude of its dissociation constant. Therefore, additional mechanisms that mediate the all-or-nothing response of EP2 could exist and might include receptor (hetero/homo) oligomerization, which are documented for other GPCRs [49] but remain to be identified for EP2 and EP4. Importantly, our results indicate that EP4 is the main producer and regulator of cAMP production at low, possibly physiological, PGE2 concentrations, whereas EP2 boosts cAMP levels only when PGE2 concentration increases above a certain threshold, as it could (locally) occur in inflamed or tumour tissues.

Interestingly, our experiments using cAMP FRET biosensor show that EP2 and EP4 both signal more strongly when stimulated selectively. This indicates that simultaneous activation of both receptors limits efficient signaling and suggests the presence of signaling crosstalk between EP2 and EP4. Since both EP2 and EP4 couple to Gα_s_, competition for downstream effectors could contribute to the attenuated cAMP response observed in the absence of receptor antagonists. Additionally, inhibitory interactions between activated receptors at the plasma membrane could attenuate the PGE2-induced cAMP response to establish an integrated signal that fine-tunes downstream effects. Although the mechanisms underlying this potential crosstalk remains to be deciphered, our results strongly indicate that the EP2 and EP4 signalling axes may be closely intertwined.

The organization of GPCR signaling has previously been linked to membrane domains and the cortical microtubule network [50]. Here, we show that an intact microtubule network is necessary for efficient signaling of both EP2 and EP4. Remarkably, several other studies show that cAMP production is dampened by intact microtubules and lipid membrane domains [50; 51; 52]. Specifically, microtubules were suggested to restrict the interactions of Gα_s_ with GPCRs and AC, limiting the efficiency of cAMP responses [51; 53]Yet, most previous research focused on adrenergic receptors, which primarily localize to lipid-raft domains [54]. By contrast, the insensitivity of EP receptors to cholesterol depletion suggests that EP2 and EP4 mainly localize in non-raft regions[55]. Moreover, the AC isoform 2, which is the AC isoform that responds most strongly to PGE2, is also located in non-raft domains, further supporting the notion that PGE2 signaling occurs outside lipid rafts and possibly explaining their differential dependence on the microtubule network that was reported for the adrenergic receptors [55]. Although a mechanistic explanation is still lacking, the different sensitivity of EP2 and EP4 to microtubule disruption is striking: whereas PGE2-EP4 signalling is partially reduced, PGE2-EP2 signalling is completely abolished by nocodazole treatment. Imaging of microtubules in combination with single-particle tracking of EP receptors could reveal the role of microtubules in PGE2 signaling. Furthermore, a detailed molecular investigation of Gα_s_ and Gα_i_ dynamics is required to accurately describe the organization and receptor-coupling of the different Gα proteins involved. The different sensitivity of EP2 and EP4 to nocodazole together with the apparently contradictory results between adrenergic and prostaglandin receptors strongly emphasizes the complexity of GPCR spatiotemporal organization and the importance of studying the regulation of a specific receptor in its endogenous settings.

Based on our experimental observations, we here present a schematic model for the cAMP responses established by EP2 and EP4. Upon selective stimulation of EP4, both Gα_s_ and Gα_i_ proteins are activated (**Figure 5A**). Active Gα_s_ proteins modulate the activity of AC, resulting in a strong cAMP response. Gα_i_ functions to fine-tune the cAMP production at low PGE2 concentrations. As EP4 is subjected to desensitization and internalization [23; 25], the elicited cAMP response subsides over time. When EP2 is selectively stimulated, only Gα_s_ controls AC activity (**Figure 5B**). The resulting cAMP response does not subside because EP2 is insensitive to receptor desensitization and internalization [23]. Disruption of the microtubule network dampens the cAMP levels induced by both EP2 and EP4, albeit with different strength, showing that microtubules play an important role in the organization of EP receptor signaling. Upon simultaneous activation of EP2 and EP4, Gα proteins are activated by both EP2 and EP4 resulting in an integrated cAMP response (**Figure 5C**). Competition between EP2 and EP4 for Gα_s_ likely reduces the signaling efficiency of individual receptors and thereby moderates final cAMP levels. Since EP4 has a higher affinity for PGE2 than EP2 [48], EP4 is the main gatekeeper of cAMP levels, especially at low PGE2 concentrations, while EP2 becomes important only at high PGE2 concentrations that will result in a strong and sustained cAMP production.

**Figure 5.**
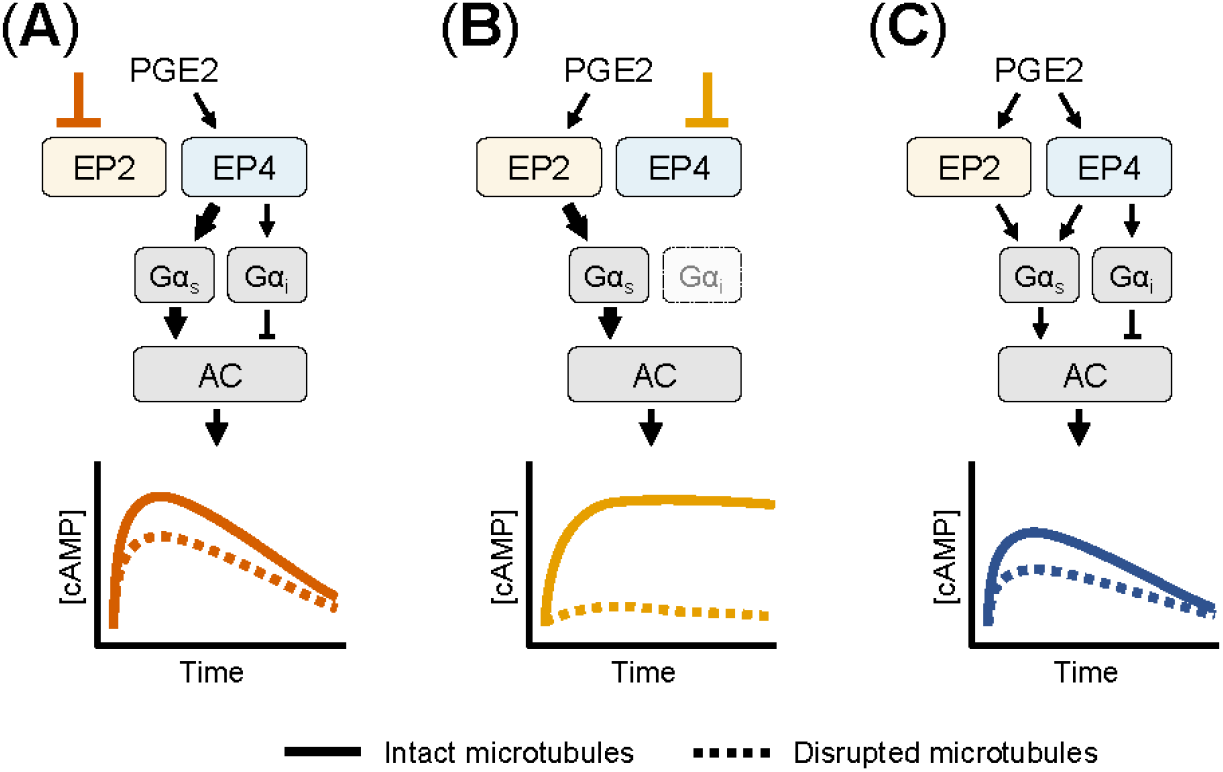
Schematic overview of the cAMP responses induced by EP2 and EP4. **(A)** When only EP4 is active, both Gα_s_ and Gα_i_ control AC activity. Gα_s_ induces a dose-dependent cAMP response that is dampened by Gα_i_. The cAMP signal subsides over time and is attenuated by microtubule disruption. **(B)** When EP2 is activated selectively, only Gα_s_ modulates AC activity. The resulting cAMP response is either weak or strong, does not subside and completely relies on an intact microtubule network. **(C)** When both EP2 and EP4 are active, competition for Gα_s_ dampens the integrated cAMP response. Signaling crosstalk between EP2 and EP4 allows the cell to respond differently to PGE2 depending on the organization and expression of EP2 and EP4.

Increased PGE2 concentrations have been reported in the tumor microenvironment of several cancer types [9; 10; 11; 12]. Since PGE2 regulates immune cell function, the selective modulation of EP receptor signaling pathways has been proven to enhance the antitumor immune response [56; 57; 58]. Further insight into the concerted action of EP2 and EP4 will be essential to efficiently control the cellular responses to PGE2.

## Author Contributions

WV, KvdD, BJ, SdK performed the experiments and analyzed the data. DSL provided analytical tools. WV, SdK, DSL and AC wrote the manuscript with input from all authors. DSL and AC superbised the entire project.

## Acknowledgements

The authors are indebted to The microscopy experiments were mostly conducted at the Radboudumc Technology Center Microscopy with the exception of the FLIM measurements that were performed at the University of Twente, Enschede, The Netherlands. The authors declare that the research was conducted in the absence of any commercial or financial relationships that could be construed as a potential conflict of interest.

## References

[1] K.L. Pierce, R.T. Premont, and R.J. Lefkowitz, Seven-transmembrane receptors. Nat.Rev.Mol.Cell Biol. 3 (2002) 639–650.

[2] W.L. Smith, Prostanoid biosynthesis and mechanisms of action. Am.J.Physiol 263 (1992) F181–F191.

[3] R.A. Coleman, W.L. Smith, and S. Narumiya, International Union of Pharmacology classification of prostanoid receptors: properties, distribution, and structure of the receptors and their subtypes. Pharmacol.Rev. 46 (1994) 205–229.

[4] D.F. Legler, M. Bruckner, E. Uetz-von Allmen, and P. Krause, Prostaglandin E2 at new glance: novel insights in functional diversity offer therapeutic chances. Int J Biochem Cell Biol 42 (2010) 198–201.

[5] S. Narumiya, Prostanoids in immunity: roles revealed by mice deficient in their receptors. Life sciences 74 (2003) 391–5.

[6] D.F. Legler, P. Krause, E. Scandella, E. Singer, and M. Groettrup, Prostaglandin E2 is generally required for human dendritic cell migration and exerts its effect via EP2 and EP4 receptors. J Immunol 176 (2006) 966–73.

[7] N. Gualde, and H. Harizi, Prostanoids and their receptors that modulate dendritic cell-mediated immunity. Immunology and cell biology 82 (2004) 353–60.

[8] S.F. van Helden, M.M. Oud, B. Joosten, N. Peterse, C.G. Figdor, and F.N. van Leeuwen, PGE2-mediated podosome loss in dendritic cells is dependent on actomyosin contraction downstream of the RhoA-Rho-kinase axis. J Cell Sci 121 (2008) 1096–106.

[9] A. Rasmuson, A. Kock, O.M. Fuskevåg, B. Kruspig, J. Simón-Santamaría, V. Gogvadze, J.I. Johnsen, P. Kogner, and B. Sveinbjörnsson, Autocrine Prostaglandin E2 Signaling Promotes Tumor Cell Survival and Proliferation in Childhood Neuroblastoma. PLOS ONE 7 (2012) e29331.

[10] B. Rigas, I.S. Goldman, and L. Levine, Altered eicosanoid levels in human colon cancer. J Lab Clin Med 122 (1993) 518–23.

[11] L.R. Howe, Inflammation and breast cancer. Cyclooxygenase/prostaglandin signaling and breast cancer. Breast Cancer Research 9 (2007) 210.

[12] M. Huang, M. Stolina, S. Sharma, J.T. Mao, L. Zhu, P.W. Miller, J. Wollman, H. Herschman, and S.M. Dubinett, Non-Small Cell Lung Cancer Cyclooxygenase-2-dependent Regulation of Cytokine Balance in Lymphocytes and Macrophages: Up-Regulation of Interleukin 10 and Down-Regulation of Interleukin 12 Production. Cancer Research 58 (1998) 1208–1216.

[13] K. Kobayashi, K. Omori, and T. Murata, Role of prostaglandins in tumor microenvironment. Cancer Metastasis Rev 37 (2018) 347–354.

[14] Y. Ma, G.V. Shurin, Z. Peiyuan, and M.R. Shurin, Dendritic cells in the cancer microenvironment. J Cancer 4 (2013) 36–44.

[15] J.S. Klarquist, and E.M. Janssen, Melanoma-infiltrating dendritic cells: Limitations and opportunities of mouse models. Oncoimmunology 1 (2012) 1584–1593.

[16] S. Kubo, H.K. Takahashi, M. Takei, H. Iwagaki, T. Yoshino, N. Tanaka, S. Mori, and M. Nishibori, E-prostanoid (EP)2/EP4 receptor-dependent maturation of human monocyte-derived dendritic cells and induction of helper T2 polarization. The Journal of pharmacology and experimental therapeutics 309 (2004) 1213–20.

[17] H. Harizi, C. Grosset, and N. Gualde, Prostaglandin E2 modulates dendritic cell function via EP2 and EP4 receptor subtypes. Journal of leukocyte biology 73 (2003) 756–63.

[18] S.F. van Helden, D.J. Krooshoop, K.C. Broers, R.A. Raymakers, C.G. Figdor, and F.N. van Leeuwen, A critical role for prostaglandin E2 in podosome dissolution and induction of high-speed migration during dendritic cell maturation. J Immunol 177 (2006) 1567–74.

[19] A. Honda, Y. Sugimoto, T. Namba, A. Watabe, A. Irie, M. Negishi, S. Narumiya, and A. Ichikawa, Cloning and expression of a cDNA for mouse prostaglandin E receptor EP2 subtype. J.Biol.Chem. 268 (1993) 7759–7762.

[20] J.W. Regan, T.J. Bailey, D.J. Pepperl, K.L. Pierce, A.M. Bogardus, J.E. Donello, C.E. Fairbairn, K.M. Kedzie, D.F. Woodward, and D.W. Gil, Cloning of a novel human prostaglandin receptor with characteristics of the pharmacologically defined EP2 subtype. Mol.Pharmacol. 46 (1994) 213–220.

[21] M. Leduc, B. Breton, C. Gales, C. Le Gouill, M. Bouvier, S. Chemtob, and N. Heveker, Functional selectivity of natural and synthetic prostaglandin EP4 receptor ligands. The Journal of pharmacology and experimental therapeutics 331 (2009) 297–307.

[22] H. Fujino, and J.W. Regan, EP4 Prostanoid Receptor Coupling to a Pertussis Toxin-Sensitive Inhibitory G Protein. Molecular pharmacology 69 (2006) 5–10.

[23] S. Desai, H. April, C. Nwaneshiudu, and B. Ashby, Comparison of agonist-induced internalization of the human EP2 and EP4 prostaglandin receptors: role of the carboxyl terminus in EP4 receptor sequestration. Mol.Pharmacol. 58 (2000) 1279–1286.

[24] R.B. Penn, R.M. Pascual, Y.M. Kim, S.J. Mundell, V.P. Krymskaya, R.A. Panettieri, Jr., and J.L. Benovic, Arrestin specificity for G protein-coupled receptors in human airway smooth muscle. The Journal of biological chemistry 276 (2001) 32648–56.

[25] S. Desai, and B. Ashby, Agonist-induced internalization and mitogen-activated protein kinase activation of the human prostaglandin EP4 receptor. FEBS letters 501 (2001) 156–60.

[26] Y. Mao, D. Sarhan, A. Steven, B. Seliger, R. Kiessling, and A. Lundqvist, Inhibition of tumor-derived prostaglandin-e2 blocks the induction of myeloid-derived suppressor cells and recovers natural killer cell activity. Clin Cancer Res 20 (2014) 4096–106.

[27] B. Thurner, C. Roder, D. Dieckmann, M. Heuer, M. Kruse, A. Glaser, P. Keikavoussi, E. Kampgen, A. Bender, and G. Schuler, Generation of large numbers of fully mature and stable dendritic cells from leukapheresis products for clinical application. J Immunol Methods 223 (1999) 1–15.

[28] I.J. de Vries, A.A. Eggert, N.M. Scharenborg, J.L. Vissers, W.J. Lesterhuis, O.C. Boerman, C.J. Punt, G.J. Adema, and C.G. Figdor, Phenotypical and functional characterization of clinical grade dendritic cells. J Immunother 25 (2002) 429–38.

[29] J.B. Klarenbeek, J. Goedhart, M.A. Hink, T.W. Gadella, and K. Jalink, A mTurquoise-based cAMP sensor for both FLIM and ratiometric read-out has improved dynamic range. PLoS.One. 6 (2011) e19170.

[30] S.K. Gibson, and A.G. Gilman, Gialpha and Gbeta subunits both define selectivity of G protein activation by alpha2-adrenergic receptors. Proc Natl Acad Sci U S A 103 (2006) 212–7.

[31] S. De Keijzer, M.B. Meddens, R. Torensma, and A. Cambi, The multiple faces of prostaglandin E2 G-protein coupled receptor signaling during the dendritic cell life cycle. Int J Mol Sci 14 (2013) 6542–55.

[32] N.E. Hubbard, S. Lee, D. Lim, and K.L. Erickson, Differential mRNA expression of prostaglandin receptor subtypes in macrophage activation. Prostaglandins, leukotrienes, and essential fatty acids 65 (2001) 287–94.

[33] M. Bünemann, M. Frank, and M.J. Lohse, Gi protein activation in intact cells involves subunit rearrangement rather than dissociation. Proceedings of the National Academy of Sciences 100 (2003) 16077–16082.

[34] N. Wang, K. Yan, and M.M. Rasenick, Tubulin binds specifically to the signal-transducing proteins, Gs alpha and Gi alpha 1. J.Biol.Chem. 265 (1990) 1239–1242.

[35] M.n. Côté, M.D. Payet, and N. Gallo-Payet, Association of αs-Subunit of the Gs Protein with Microfilaments and Microtubules: Implication during Adrenocorticotropin Stimulation in Rat Adrenal Glomerulosa Cells1. Endocrinology 138 (1997) 69–78.

[36] T. Sarma, T. Voyno-Yasenetskaya, T.J. Hope, and M.M. Rasenick, Heterotrimeric G-proteins associate with microtubules during differentiation in PC12 pheochromocytoma cells. Faseb J 17 (2003) 848–59.

[37] E. Scandella, Y. Men, S. Gillessen, R. Forster, and M. Groettrup, Prostaglandin E2 is a key factor for CCR7 surface expression and migration of monocyte-derived dendritic cells. Blood 100 (2002) 1354–61.

[38] K. Kabashima, D. Sakata, M. Nagamachi, Y. Miyachi, K. Inaba, and S. Narumiya, Prostaglandin E2-EP4 signaling initiates skin immune responses by promoting migration and maturation of Langerhans cells. Nature medicine 9 (2003) 744–9.

[39] G. Flórez-Grau, R. Cabezón, K.J.E. Borgman, C. España, J.J. Lozano, M.F. Garcia-Parajo, and D. Benítez-Ribas, Up-regulation of EP2 and EP3 receptors in human tolerogenic dendritic cells boosts the immunosuppressive activity of PGE2. Journal of leukocyte biology 102 (2017) 881–895.

[40] N.J. Poloso, P. Urquhart, A. Nicolaou, J. Wang, and D.F. Woodward, PGE(2) differentially regulates monocyte-derived dendritic cell cytokine responses depending on receptor usage (EP(2)/EP(4)). Molecular immunology 54 (2013) 284–295.

[41] C. Yao, D. Sakata, Y. Esaki, Y. Li, T. Matsuoka, K. Kuroiwa, Y. Sugimoto, and S. Narumiya, Prostaglandin E2–EP4 signaling promotes immune inflammation through TH1 cell differentiation and TH17 cell expansion. Nature medicine 15 (2009) 633–640.

[42] H. Fujino, K.A. West, and J.W. Regan, Phosphorylation of glycogen synthase kinase-3 and stimulation of T-cell factor signaling following activation of EP2 and EP4 prostanoid receptors by prostaglandin E-2. Journal of Biological Chemistry 277 (2002) 2614–2619.

[43] H. Fujino, W. Xu, and J.W. Regan, Prostaglandin E2 induced functional expression of early growth response factor-1 by EP4, but not EP2, prostanoid receptors via the phosphatidylinositol 3-kinase and extracellular signal-regulated kinases. J.Biol.Chem. 278 (2003) 12151–12156.

[44] H. Fujino, S. Salvi, and J.W. Regan, Differential regulation of phosphorylation of the cAMP response element-binding protein after activation of EP2 and EP4 prostanoid receptors by prostaglandin E2. Molecular pharmacology 68 (2005) 251–9.

[45] F.F. Hamdan, M.D. Rochdi, B. Breton, D. Fessart, D.E. Michaud, P.G. Charest, S.A. Laporte, and M. Bouvier, Unraveling G Protein-coupled Receptor Endocytosis Pathways Using Real-time Monitoring of Agonist-promoted Interaction between β-Arrestins and AP-2. Journal of Biological Chemistry 282 (2007) 29089–29100.

[46] A. Bondar, and J. Lazar, G protein Gi1 exhibits basal coupling but not preassembly with G protein-coupled receptors. Journal of Biological Chemistry (2017).

[47] T. Sungkaworn, M.-L. Jobin, K. Burnecki, A. Weron, M.J. Lohse, and D. Calebiro, Single-molecule imaging reveals receptor–G protein interactions at cell surface hot spots. Nature 550 (2017) 543–547.

[48] M. Abramovitz, M. Adam, Y. Boie, M. Carriere, D. Denis, C. Godbout, S. Lamontagne, C. Rochette, N. Sawyer, N.M. Tremblay, M. Belley, M. Gallant, C. Dufresne, Y. Gareau, R. Ruel, H. Juteau, M. Labelle, N. Ouimet, and K.M. Metters, The utilization of recombinant prostanoid receptors to determine the affinities and selectivities of prostaglandins and related analogs. Biochimica et biophysica acta 1483 (2000) 285–93.

[49] R. Sleno, and T.E. Hebert, The Dynamics of GPCR Oligomerization and Their Functional Consequences. Int Rev Cell Mol Biol 338 (2018) 141–171.

[50] B.P. Head, H.H. Patel, D.M. Roth, F. Murray, J.S. Swaney, I.R. Niesman, M.G. Farquhar, and P.A. Insel, Microtubules and actin microfilaments regulate lipid raft/caveolae localization of adenylyl cyclase signaling components. The Journal of biological chemistry 281 (2006) 26391–9.

[51] S.M. Pontier, Y. Percherancier, S. Galandrin, A. Breit, C. Galés, and M. Bouvier, Cholesterol-dependent Separation of the β2-Adrenergic Receptor from Its Partners Determines Signaling Efficacy: INSIGHT INTO NANOSCALE ORGANIZATION OF SIGNAL TRANSDUCTION. Journal of Biological Chemistry 283 (2008) 24659–24672.

[52] J.A. Allen, J.Z. Yu, R.H. Dave, A. Bhatnagar, B.L. Roth, and M.M. Rasenick, Caveolin-1 and Lipid Microdomains Regulate G_s_ Trafficking and Attenuate G_s_/Adenylyl Cyclase Signaling. Molecular pharmacology 76 (2009) 1082–1093.

[53] A.H. Czysz, J.M. Schappi, and M.M. Rasenick, Lateral Diffusion of Gα_s_ in the Plasma Membrane Is Decreased after Chronic but not Acute Antidepressant Treatment: Role of Lipid Raft and Non-Raft Membrane Microdomains. Neuropsychopharmacology 40 (2015) 766–773.

[54] S.R. Agarwal, P.-C. Yang, M. Rice, C.A. Singer, V.O. Nikolaev, M.J. Lohse, C.E. Clancy, and R.D. Harvey, Role of Membrane Microdomains in Compartmentation of cAMP Signaling. PLOS ONE 9 (2014) e95835.

[55] A.S. Bogard, P. Adris, and R.S. Ostrom, Adenylyl Cyclase 2 Selectively Couples to E Prostanoid Type 2 Receptors, Whereas Adenylyl Cyclase 3 Is Not Receptor-Regulated in Airway Smooth Muscle. Journal of Pharmacology and Experimental Therapeutics 342 (2012) 586–595.

[56] X. Ma, D. Holt, N. Kundu, J. Reader, O. Goloubeva, Y. Take, and A.M. Fulton, A prostaglandin E (PGE) receptor EP4 antagonist protects natural killer cells from PGE-mediated immunosuppression and inhibits breast cancer metastasis. Oncoimmunology 2 (2013) e22647.

[57] M. Majumder, X. Xin, L. Liu, G.V. Girish, and P.K. Lala, Prostaglandin E2 receptor EP4 as the common target on cancer cells and macrophages to abolish angiogenesis, lymphangiogenesis, metastasis, and stem-like cell functions. Cancer Sci 105 (2014) 1142–51.

[58] D.I. Albu, Z. Wang, K.C. Huang, J. Wu, N. Twine, S. Leacu, C. Ingersoll, L. Parent, W. Lee, D. Liu, R. Wright-Michaud, N. Kumar, G. Kuznetsov, Q. Chen, W. Zheng, K. Nomoto, M. Woodall- Jappe, and X. Bao, EP4 Antagonism by E7046 diminishes Myeloid immunosuppression and synergizes with Treg-reducing IL-2-Diphtheria toxin fusion protein in restoring anti-tumor immunity. Oncoimmunology 6 (2017) e1338239.

